# Chronic Aryl Hydrocarbon Receptor Activity Phenocopies Smoking-induced Skeletal Muscle Impairment

**DOI:** 10.1101/2021.05.05.441943

**Authors:** Trace Thome, Kayla Miguez, Alexander Willms, Angela R. de Souza, Vijayendran Chandran, Sarah S. Burke, Yana Goddard, Carolyn Baglole, Maria-Eleni Anagnostou, Jean Bourbeau, R. Thomas Jagoe, Jose Morais, Tanja Taivassalo, Terence Ryan, Russell T. Hepple

**Affiliations:** Department of Applied Physiology & Kinesiology, University of Florida, USA; Department of Kinesiology and Physical Education, McGill University, CANADA; Research Institute of the McGill University Health Center, McGill University, CANADA; Department of Pediatrics, University of Florida, USA; Department of Physical Therapy, University of Florida, USA; Department of Medicine, University of Florida, USA; Department of Physiology & Functional Genomics, University of Florida, USA

**Keywords:** smoking, cachexia, sarcopenia, neuromuscular junction

## Abstract

**Background:** COPD patients exhibit skeletal muscle atrophy, denervation, and reduced mitochondrial oxidative capacity. Whilst chronic tobacco smoke exposure is implicated in COPD muscle impairment, the mechanisms involved are ambiguous. The aryl hydrocarbon receptor (AHR) is a ligand-activated transcription factor that activates detoxifying pathways with numerous exogenous ligands, including tobacco smoke. Whereas transient AHR activation is adaptive, chronic activation can be toxic. On this basis, we tested the hypothesis that chronic smoke-induced AHR activation causes adverse muscle impact.

**Methods:** We used clinical patient muscle samples, and *in vitro* (C2C12 myotubes) and *in vivo* models (mouse), to perform gene expression, mitochondrial function, muscle and neuromuscular junction morphology, and genetic manipulations (adeno-associated virus-mediated gene transfer).

**Results:** 16 weeks tobacco smoke exposure in mice caused: muscle atrophy, neuromuscular junction degeneration, and reduced oxidative capacity. Similarly, smoke exposure reprogrammed the muscle transcriptome, with down-regulation of mitochondrial and neuromuscular junction genes. In mouse and human patient specimens, smoke exposure increased muscle AHR signaling. Mechanistically, experiments in cultured myotubes demonstrated that smoke condensate activated the AHR, caused mitochondrial impairments, and induced an AHR-dependent myotube atrophy. Finally, to isolate the role of AHR activity, expression of a constitutively active AHR mutant without smoke exposure caused atrophy and mitochondrial impairments in cultured myotubes, and muscle atrophy and neuromuscular junction degeneration in mice.

**Conclusions:** These results establish that chronic AHR activity, as occurs in smokers, phenocopies the atrophy, mitochondrial impairment and neuromuscular junction degeneration caused by chronic tobacco smoke exposure.

## INTRODUCTION

Smoking is a major risk factor for several diseases, including chronic obstructive pulmonary disease (COPD), cardiovascular disease, and many cancers [41]. Interestingly, patients with smoking-related disease often exhibit skeletal muscle atrophy, where atrophy reduces quality of life, increases hospitalizations, and increases the risk of death [20, 24]. Whereas the role of chronic tobacco smoking in developing COPD is well-established [23], and tobacco smoke is known to have dose-dependent adverse impact on skeletal muscle [11], the cellular mechanisms by which smoking contributes to the muscle impairment often seen in COPD patients are poorly understood.

Interestingly, numerous constituents of tobacco smoke activate the aryl hydrocarbon receptor (AHR) [19], a ligand-activated transcription factor best known for mediating the toxic effects of 2,3,7,8-tetrachlorodibenzodioxin (dioxin) [27]. Activation of the AHR regulates cytochrome P450 enzymes such as Cyp1A1 and Cyp1B1, as well as antioxidant pathways that include NAD[P]H quinone dehydrogenase and sulfiredoxin [27, 38]. Whereas the AHR plays an important role in normal development [30], chronic AHR activation can have pathological consequences due to mitochondrial-mediated oxidative stress [1] for various organ systems [28], including the brain and nervous system, reproductive organs, heart, liver, and immune system [6]. Notably, although the impact of chronic AHR activity in muscle has not been directly studied, chronic exposure to environmental AHR agonists, including dioxin [25], tobacco smoke [47] and pesticides [48], is associated with muscle atrophy.

On the basis of the above, we sought to determine whether AHR signaling is elevated in skeletal muscle of smokers, and test the hypothesis that chronic AHR activation, with or without tobacco smoke exposure, induces adverse muscle affect. Consistent with this hypothesis, genes indicative of AHR activation are elevated in muscle of smokers and tobacco smoke condensate in cultured myotubes causes muscle atrophy that is prevented by antagonizing AHR signaling. Furthermore, knock-in of a constitutively active mutant of the AHR causes myotube atrophy and impaired mitochondrial function *in vitro*, and in mice causes muscle atrophy and neuromuscular junction degeneration; changes that are similar to those induced by chronic smoke exposure. We conclude that chronic smoking-induced AHR activation has an important role in causing adverse muscle impacts, such as those commonly seen in patients with COPD.

## Methods

### Smoking Mouse Model

To model the impact of chronic tobacco smoke exposure on skeletal muscle, we used a smoking mouse model that we have previously described [17]. The first set of experiments were designed to establish the impact of chronic smoke exposure on muscle mass, muscle fiber size, mitochondrial function, and extent of neuromuscular junction denervation in limb muscle using male C57BL/6 mice (n=24) obtained from the in-house colony at the Research Institute of the McGill University Health Center (RI-MUHC). The second set of experiments were designed to provide information about the impact of chronic smoke exposure on motor axons, and extend observations to breathing muscle, using male C57BL/6-elite mice (n=15) purchased from Charles River. In this latter respect, the C57BL/6-elite mice differ from C57BL/6 mice in that the former are raised under conditions where specific infectious agents are excluded from the colony to maintain a virus antigen free condition. In both sets of experiments, mice were maintained on a 12:12 dark/light schedule in the RI-MUHC vivarium, housed 1-4 per cage and provided with water and food *ad libitum*. All procedures in mice were done in compliance with the regulations of the Canadian Council on Animal Care and with prior approval from the McGill University Animal Care Committee (protocol 5933).

Tobacco smoke exposures were done using a SCIREQ InExpose System (SCIREQ, Montreal, QC, CANADA) and used a smoking protocol approved by the Federal Trade Commission (1 puff per min per cigarette, where each puff was 2 s in duration and 35 ml in volume). Mice in the smoke exposed group were exposed for 60 min twice per d, 5 d per week, for 16 weeks (first set of experiments: n=13; second set of experiments: n=8). Mice in the Air exposure group (first set of experiments: n=11; second set of experiments: n=7) were brought to the same room as the smoke exposure system but remained in their cages. Particulate matter density during smoke exposures were maintained between 4.0 and 6.0 mg·m^-3^·min^-1^ (measured by MicroDust Pro, Casella, Buffalo NY, USA). The sample size used in each assay is indicated below.

### Tissue Harvest

Mice in the first set of experiments were sacrificed by CO_2_ asphyxiation followed by cervical dislocation, 24 h after their last smoke or Air exposure. The soleus (Sol), plantaris (Plan), gastrocnemius (Gas), and diaphragm (Dia) muscles were rapidly removed, dissected free of connective tissue and fat, and weighed. The tibialis anterior (TA) in 5 animals per group (Air, TS) was also removed and prepared for neuromuscular junction labeling as previously described [17], and is briefly described below. The central tendon and crural Dia were separated from the whole Dia and discarded. A portion of the remaining costal Dia was prepared for mitochondrial function assays (see below). Similarly, the right Sol was used in mitochondrial function assays, whereas the left Sol was mounted for histology. The right and left plantaris muscles were flash-frozen in liquid N_2_ immediately following weighing to be used in transcriptional analyses (see Quantseq, below).

Mice in the second set of experiments were sacrificed by cervical dislocation after anesthetizing with ketamine (1 ml per g body mass), 48 h after their last smoke or Air exposure. The Dia was removed and prepared as above to isolate the crural Dia. The crural Dia was then prepared for neuromuscular junction labeling as detailed below.

### Histology

For mice in the first set of experiments, the entire left Sol muscle was mounted in Cryomatrix embedding resin (Thermo Scientific, USA) and frozen in liquid isopentane that had been chilled to the point of freezing in liquid N_2_, and was then stored at −80°C until processed. Muscles were cut on a cryostat (−18°C) to 10-μM sections, labeled for laminin to demarcate muscle fiber borders, and imaged on a Zeiss Axio Imager M2 fluorescence microscope. Briefly, 10 μm thick muscle cross-sections were rehydrated with PBS (pH 7.2) and then blocked using goat serum (10% in PBS) by incubating for 1 h at room temperature. The sections were then incubated overnight at 4°C with polyclonal anti-laminin (1:1000; ThermoFisher Scientific) primary antibody diluted in blocking solution. The muscle sections were then washed three times in PBS before being incubated for 1 h in the dark at room temperature with Alexa Fluor 488 IgG secondary antibody (A-11008, 1:500; ThermoFisher Scientific) diluted in blocking solution. Muscle cross-sections were then washed three times in PBS, and coverslips were applied to slides using Prolong Gold (ThermoFisher Scientific; P36930) as mounting medium. Slides were imaged using a Zeiss Axio Imager (Zeiss, Germany). Fiber size was assessed using Image J on tiled images on an average of 277 ± 33 fibers per muscle using a systematic sampling strategy to provide unbiased estimates (n=6 mice per group).

### Neuromuscular Junction Morphology

We have reported some indices of neuromuscular junction morphology from the TA muscle of the animals studied in the first set of experiments in our previous publication [17]. Briefly, TA muscles were washed (3 x 5 min) in PBS and fixed overnight at 4°C in 4% paraformaldehyde. The deep (oxidative) portion of the fixed muscles were then mechanically separated into smaller bundles. Muscle bundles were incubated overnight at 4°C in a blocking solution consisting of 5% normal goat serum, 5% BSA, and 2% Triton X-100 in PBS. Pre-synaptic motoneuron terminals were then labeled by incubating bundles overnight at 4°C in the same blocking solution to which had been added a mouse anti-synaptophysin antibody (AB8049 Abcam, USA; 1:25 dilution). The following morning bundles were washed (5 x 30 min) in a blocking solution containing 5% normal goat serum and 5% BSA in PBS, and then incubated overnight at 4°C with AF594-conjugated goat anti-mouse IgG1 secondary antibody (A21125, Invitrogen, USA; 1:500 dilution) and AF488-conjugated α-bungarotoxin (B13422, Life Technologies, USA; 1:500 dilution) to visualize the synaptophysin labeling and the AChR cluster on the muscle fiber, respectively. After incubation, muscle bundles were washed (5 x 45 min) in the blocking solution, and mounted onto slides with ProLong Gold Antifade Mountant (P36930, Life Technologies, USA).

For mice in the second set of experiments, a portion of the costal Dia was prepared as the TA above, with the exception that in addition to labeling for synaptophysin and α-bungarotoxin, we also labeled for neurofilament 200 (NF200) to label the motor axons. Briefly, after being harvested the costal Dia was immediately washed in 1 x PBS solution for 20 min. Using 2% paraformaldehyde (PFA) solution, Dia muscles were fixed for 4 h at room temperature, washed (1 x 20 min, followed by 3 x 5 min), and stored in 1 x PBS solution at 4°C for 1 to 36 h prior to permit batch processing for neuromuscular junction labeling. The length of this storage time was randomized amongst the two groups. Costal Dia was then cut widthwise into smaller pieces and incubated overnight at 4°C in blocking solution as in TA (above). Muscles were then incubated in blocking solution containing rabbit anti-heavy NF200 primary antibody (N4142 Sigma, Germany; 1:200 dilution) and anti-synaptophysin antibody (1:25 dilution as above). The following morning bundles were washed (5 x 60 min) in a blocking solution containing 5% normal goat serum and 5% BSA in PBS, and then incubated overnight at 4°C with AF594-conjugated goat anti-mouse IgG1 secondary antibody (A21125, ThermoFisher, USA; 1:500 dilution), Cy5 goat anti-rabbit IgG (H&L) (A10523, ThermoFisher, USA; 1:200) and AF488-conjugated α-bungarotoxin (B13422, Life Technologies, USA; 1:500 dilution). Muscle bundles were washed (5 x 60 min) in the blocking solution, stored in blocking solution overnight at 4°C, and then mounted onto slides with ProLong Gold as above.

### Neuromuscular Junction Imaging

Both TA (mice in first set of experiments) and Dia (mice in second set of experiments) muscles labeled for neuromuscular junction structures were imaged using an LSM880 confocal microscope (Carl Zeiss, Germany) with a 63x oil-immersion objective by generating image stacks (optical slice thickness = 1.0 μm). Image stacks were then analyzed as maximum intensity projections using Image J. For the first set of experiments where only the AChRs and motoneuron terminals were labeled in TA muscles, analyses focused on identifying the fraction of neuromuscular junctions lacking motoneuron terminals (=no synaptophysin opposing the α-bungarotoxin-labeled AChRs). For the second set of experiments where we labeled the motor axons (NF200), terminals (synaptophysin) and AChRs (α-bungarotoxin) in Dia muscle, our measurements were based upon an analysis scheme developed by Jones and colleagues [15] that assessed structural features of the motor axons, the motoneuron terminals, and the AChR clusters.

### Mitochondrial Function and Content

As noted in the description of the Tissue Harvest (above), for animals in the first set of experiments the right Sol and a portion of the Dia were used in mitochondrial function assays, using methods we have described previously [42]. Briefly, freshly dissected muscle was immediately placed in ice-cold stabilizing Solution A (2.77mM CaK_2_EGTA, 7.23mM K_2_EGTA, 6.56 MgCl_2_, 0.5mM dithiothreitol, 50mM K-MES, 20mM imidazol, 20mM taurine, 5.3mM Na_2_ATP, 15mM phosphocreatine). The muscles were then dissected into smaller bundles of muscle fibers in Solution A on ice under a stereomicroscope, before being placed in 6 ml of Solution A supplemented with 200 μL of a 50 μg·ml^-1^ saponin solution and incubated on ice on a shaking platform for 30 min. Fiber bundles were then washed (3 x 5 min) in Solution B (2.77 mM CaK_2_EGTA, 7.23 mM K_2_EGTA, 1.38 mM MgCl_2_, 0.5 mM dithiothreitol, 100 mM K-MES, 20 mM imidazol, 20 mM Taurine, 3 mM K_2_HPO_4_) supplemented with 2 mg x ml^-1^ of bovine serum albumin (BSA). Washed fiber bundles weighing 3-6 mg were then placed in each of the two chambers of an Oroboros Oxygraph O2K (Oroboros, Austria), in Solution B maintained at 37°C. The rate of respiration was then measured in response to the following substrate-inhibitor protocol: (1) glutamate (10 mM) and malate (5 mM), (2) ADP (5 mM), (3) succinate (20 mM), (4) cytochrome C (10 μM), (5) antimycin A (10 μM), and (6) ascorbate (12.5 mM) and the artificial electron donor N,N,N·,N’-tetramethyl-p-phenylenediamine dihydrochloride (TMPD; 1.25 mM).

Following the mitochondrial respiration assessment, remaining Sol and Dia fiber bundles were blotted dry using a Kimwipe, and frozen in liquid N_2_ for use in determining the amount of the mitochondrial outer membrane protein, voltage dependent anion channel (VDAC). Briefly, approximately 10 mg of muscle was homogenized in a MM400 robot homogenizer (Retsch, Germany) with 10 x weight per volume of extraction buffer (50 mM Tris base, 150 mM NaCl, 1% Triton X-100, 0.5% sodium deoxycolate, 0.1% sodium dodecyl sulfate, and 10 μl·ml^-1^ Protease Inhibitor Cocktail (Sigma, USA)). Following 2 h gentle agitation at 4°C, samples were centrifuged at 12,000 g at 4°C for 20 min. The supernatant was then removed and samples were diluted in 4 x Laemli buffer to a final protein concentration of 2 μg·ml^-1^ before boiling for 5 min at 95°C. We then performed immunoblotting by loading 20 μg of tissue protein onto a 12% acrylamide gel, electrophoresed by SDS-PAGE and then transferred to polyvinylidene fluoride membranes (Life Sciences) blocked for 1 h with 5% (w/v) semi-skimmed milk at room temperature. Gels were then probed overnight at 4°C with mouse monoclonal anti-VDAC (ab14734, Abcam; 1:1000). Ponceau staining was done to permit normalizing to protein loading. After washing, membranes were incubated with HRP-conjugated secondary antibody (Abcam; diluted in 5% milk) at room temperature for 1 h. Protein bands were detected by SuperSignal™ West Pico Chemiluminescent Substrate (Thermo Scientific, USA) and imaged with a G-Box chem imaging system.

### Quantseq Analysis

Flash frozen plantaris muscles from mice studied in the first set of experiments were used in analyses of the muscle transcriptome response to chronic smoke exposure. Frozen muscle tissue (30 mg) was homogenized using the Fisherbrand Bead Mill 4 and total RNA was extracted from homogenates using the RNeasy Tissue Mini Kit (Qiagen), according to manufacturer’s instructions. RNA concentration and purity (A260/A280 ratios >1.8) were assessed using a spectrophotometer. RNA was also tested for suitable mass (RiboGreen) and integrity (Agilent TapeStation), reverse transcribed to complementary DNA (Lexogen QuantSeq 3’ FWD), and sequenced on a HiSeq 4000 instrument (Illumina) in the UCLA Neuroscience Genomics Core laboratory, following the manufacturers’ standard protocols. Sequencing targeted mean 7 million 65-nt single-stranded reads per sample, which were mapped to the mouse transcriptome and quantified as transcripts per million mapped reads using the STAR aligner. All sequencing data were uploaded to Illumina’s BaseSpace in real-time for downstream analysis of quality control. Raw Illumina (fastq.gz) sequencing files were downloaded from BaseSpace, and uploaded to Bluebee’s genomics analysis platform (https://www.bluebee.com) to align reads against the mouse genome. After combining treatment replicate files, a DESeq2 application within Bluebee (Lexogen Quantseq DE 1.2) was used to identify significant treatment-related effects on transcript abundance (relative to controls) based on a false discovery rate (FDR) p-adjusted value <0.1.

### Construction of co-expression networks

A weighted signed gene co-expression network was constructed using the normalized dataset to identify groups of genes (modules) associated with chronic smoke exposure following a previously described algorithm[31, 50]. Briefly, we first computed the Pearson correlation between each pair of selected genes yielding a similarity (correlation) matrix. Next, the adjacency matrix was calculated by raising the absolute values of the correlation matrix to a power (β) as described previously[50]. The parameter β was chosen by using the scale-free topology criterion[50], such that the resulting network connectivity distribution best approximated scale-free topology. The adjacency matrix was then used to define a measure of node dissimilarity, based on the topological overlap matrix, a biologically meaningful measure of node similarity[50]. Next, the genes were hierarchically clustered using the distance measure and modules were determined by choosing a height cutoff for the resulting dendrogram by using a dynamic tree-cutting algorithm[50]. Utilizing this network analysis, we identified modules (groups of genes) differentially expressed across different sample data sets after chronic smoke exposure and calculated the first principal component of gene expression in each module (module eigengene). Next, we correlated the module eigengenes with chronic smoke exposure treatment to select modules for functional validation. Gene ontology and pathway enrichment analysis was performed using the DAVID platform[14] (DAVID, https://david.ncifcrf.gov/). A list of differentially regulated transcripts for a given modules were utilized for enrichment analyses. Datasets generated and analyzed in this study are available at Gene Expression Omnibus, GEO accession: GSE151099.

### Human Muscle Biopsies

Muscle biopsies were obtained by the Bergstrom method from two clinical populations with a high smoking incidence (COPD and Critical Limb Ischemia [CLI]). Vastus lateralis muscle biopsies were obtained from 9 ambulatory male COPD patients (aged 58-77 y) and 6 healthy adult controls (aged 20-72 y) from our previous report [17]. As previously reported [17], these biopsies were obtained with approval from the Institutional Review Board for human studies at the Montreal Chest Institute (Montreal, Canada; #BMC-08-026) and Research Institute of the McGill University Health Center (#BMC-06-015). All subjects provided written informed consent. Of the 9 COPD patients, 6 were former smokers (age: 65.5 ± 6.8 y; FEV-1: 1.08 ± 0.33 L; PaO_2_: 74.7 ± 15.8 mmHg; PaCO_2_: 41.7 ± 5.1 mmHg) and 3 were current smokers (age: 63.7 ± 4.0 y; FEV-1: 1.16 ± 0.43 L; PaO_2_: 68.3 ± 8.1 mmHg; PaCO_2_: 42.0 ± 6.2 mmHg). Gastrocnemius muscle biopsies were obtained from 18 patients with CLI undergoing limb amputation classified as either current smokers (n=8; 58.4 ± 7.0 y; Ankle brachial index: 0.59 ± 0.15) or non-smokers (never smoked or quit more than 2 years prior to tissue acquisition; n=10; 59.7 ± 8.5 y; Ankle brachial index: 0.63 ± 0.23). Biopsies were acquired with approval by the institutional review board at the University of Florida (Gainesville, FL, USA; IRB201802025). All participants were fully informed about the research and informed consent was obtained. In both cases, 20 mg pieces of muscle were fast-frozen in liquid N_2_ for subsequent mRNA analysis (described below). All human studies were carried out according to the Declaration of Helsinki.

### Muscle Cell Culture

Murine skeletal C2C12 myoblasts were obtained from ATCC, USA (CRL-1772) and cultured in Dulbecco’s Modified Eagle Medium + GlutaMAX (DMEM: Cat. No. 10569, Gibco, USA) supplemented with 10% Fetal Bovine Serum (FBS: Cat. No. 97068, VWR, USA) and 1% Penicillin/Streptomycin (Cat. No. 15140, Gibco, USA) at standard culture conditions (37°C in 5% CO_2_). All cell experiments were performed with 3-4 biologically independent cell samples. To generate mature myotubes, confluent myoblast cultures were subjected to serum withdrawal by switching DMEM medium from 10% FBS to 2% adult horse serum. This differentiation medium was changed every 24 hours for six days to form mature myotubes. For experiments involving TS extract (TSE) treatment, TSE was obtained from Murty Pharmaceuticals (USA) and used at a final concentration of 0.02%. TSC was prepared by smoking University of Kentucky’s 3R4F Standard Research Cigarettes on an FTC Smoke Machine. The Total Particulate Matter (TPM) on the filter was calculated by the weight gain of the filter. From the TPM, the amount of DMSO to be used for extraction to prepare a 4% (40 mg/mL) solution is calculated. The condensate is extracted with DMSO by soaking and sonication. For experiments involving genetic knockdown (shRNA) of the AHR, myotubes were transfected 48h prior to TSE exposure. For experiments involving chemical AHR antagonism with 25 μM resveratrol or 1 μM CH223191, chemical antagonists were provided to cells three hours before TSE and remained throughout the entire treatment period (24h).

### Measurement of Myotube Viability

Cell viability was assessed by incubating live myotubes with 10 μM Ethidium Homodimer-1 (EtHD-1, Cat. No. 46043, Millipore Sigma, USA) and 1 μM Calcein AM (to label live myotubes). EtHD-1 is a cell-impermeant viability indicator that is strongly fluorescent when bound to DNA. Myotubes treated with 0.25% Triton X-100 in HBSS to permeabilize cell membranes were used as a positive control. EtHD-1 positive nuclei were quantified and expressed as a percentage of the Triton X-100 treated control cells using custom batch processing routines in Cell Profiler (The Broad Institute, USA).

### Myotube Respiration and ROS Production

Myotube respiration and superoxide production were performed as previously described [44]. For respiration measurements, high resolution respirometry was performed using an Oroboros Oxygraph-2K (Austria). Myotubes were gently rinsed with PBS, trypsinized to detach from well plates, and collected using centrifugation at 500 g. The resulting myotubes were resuspended in 2.5ml of Buffer Z (105 mM K-MES, 30 mM KCl, 1 mM EGTA, 10 mM K2HPO4, 5 mM MgCl2-6H2O, 2.5 mg/ml BSA, pH 7.1) supplemented with glucose (10mM) and pyruvate (5mM). An aliquot of the myotube suspension was used to measure protein content to normalize respiration rates accordingly. Cells were loaded into the oxygraph chamber (O2K, Oroboros, Austria) and respiration was measured at 37°C. Basal oxygen consumption (*J*O_2_) was measured in intact myotubes followed by a titration of carbonyl cyanide 4-(trifluoromethoxy)phenylhydrazone (FCCP; 250nM-1.5μM) to stimulate maximal uncoupled respiration, followed by the addition of rotenone (0.01 mM) and antimycin A (0.005 mM) to account for non-mitochondrial oxygen consumption.

To assess mitochondrial superoxide production, live myotubes were washed twice with Hanks balanced salt solution (HBSS) and incubated with 500nM MitoSOX for 15 min in HBSS prior to imaging. Live images at 20x were captured using and Evos FL2 Auto fluorescent microscope using automated capture to avoid any human bias. MitoSOX positive area was calculated using automated analysis routines created in Cell Profiler (Broad Institute, USA). All processing procedures were performed uniformly over the entire set of images using batch processing modes to avoid any human bias.

### Measurement of Myotube Area

Myotube area was measured as previously described [4, 26]. Treated myotubes were gently washed with PBS, fixed with 1:1 methanol:acetone for ten minutes at −20°C, left to air dry for 10 min, and incubated with primary antibody against sarcomeric myosin (MF 20 was deposited to the DSHB by Fischman, D.A. (Product MF 20; DSHB Hybridoma Bank, USA) at 1:25 in blocking solution (PBS + 5% goat serum + 1% BSA) for one hour at 37°C. Cells were then washed 3x in PBS, followed by incubation with 1:250 secondary antibody (AlexaFluor594, mouse IgG2b; ThermoFisher, USA) for one hour at 37°C. Cells were imaged using automated capture routines on an Evos FL Auto 2 inverted fluorescent microscope (ThermoFisher, USA) and analyzed using custom written routines in CellProfiler (Broad Institute, USA) to assess MF20+ area (myotube area). All processing procedures were performed uniformly over the entire set of images using batch processing modes to avoid any human bias.

### Plasmid Construction and AAV Production/Delivery

AAV backbones were obtained from Cell Biolabs, USA (Cat. No. VPK-411-DJ). The AAV-CMV-GFP plasmid was developed by inserting a CMV promoter and GFP (ZsGreen1) into a promoterless AAV vector (Cat. No. VPK-411-DJ; Cell BioLabs, USA) using In-Fusion Cloning (Cat. No. 638911; Takare Bio, USA). To generate a constitutively active AHR (CAAHR) vector, the mouse AHR coding sequence was PCR amplified from genomic DNA obtained from a C57BL6J mouse such that the ligand binding domain (amino acids 288-421) was deleted as previously described for the murine AHR [2], and subsequently cloned into the pAAV-CMV vector using In-Fusion cloning. To generate inducible shRNA vectors to knockdown the murine AHR, the pLKO.1 cloning plasmid was obtained from Addgene, USA (Cat. No. 10878) [29]. The shRNA cassette from the pLKO.1 plasmid was PCR amplified and subsequently inserted into a promoterless AAV plasmid (CellBio Labs, USA). A knockdown sequence for the murine AHR (TRCN0000055409 was obtained from Millipore-Sigma) and inserted into the resulting pAAV-shRNA cloning vector using AgeI and EcoRI restriction sites. For muscle cell culture experiments, plasmids were transfected into C2C12 cells using Xfect reagent (Cat. No. 631324; Takara Bio, USA) according to manufacturer instructions.

AAV-DJ were produced using triple-transfection of HEK293T cells using the DJ-packaging kit from Cell Biolabs, USA (Cat. No. VPK-411-DJ). AAV purification was performed ~72h after triple-transfection using purification kits from Takara Bio, USA (Cat. No. 6666) according to manufacturer instructions. Purified AAVs were titered using a qPCR based kit (Cat. No. 6233; Takara Bio, USA). The TA muscle of each mouse received a 5 x 10^10^ vg of either AAV-GFP or AAV-CAAHR using several small volume (~7-8 μl) injections to ensure adequate spatial distribution across the muscle.

### Real-time qPCR

We performed real-time qPCR to assess the impact of TS exposure on components of the AHR signaling pathway in C2C12 muscle cell culture and human muscle samples from CLI and COPD patients. Total RNA was extracted from treated C2C12 myotube cultures using Trizol-Phenol Reagent (Invitrogen; Cat. No. 15596026) as described by manufacturer’s instructions. RNA quantity and quality was assessed using UV-spectroscopy (ThermoFisher Scientific; Nanodrop 2000). cDNA was generated from 500 ng RNA using Superscript IV (ThermoFisher; Cat. No. 18091200) according to manufacture directions. Real-time PCR (RT-PCR) was performed on a Quantstudio 3 (ThermoFisher Scientific) using Taqman Fast Advanced Master mix (ThermoFisher Scientific; Cat. No. 4444963) and Taqman FAM-labeled probes for AHR (ThermoFisher Scientific; Mm00478932_m1), CYP1A1 (ThermoFisher Scientific; Mm00487218_m1), or CYP1B1 (ThermoFisher Scientific; Mm00487229_m1) multiplexed with VIC-labeled probe for 18S (ThermoFisher Scientific; Hs03003631_g1). Relative gene expression was calculated using 2^-ΔΔCT^ from the respective control group for each experiment.

For the human patient samples, ~20 mg pieces of snap-frozen muscle tissue were placed into an eppendorf tube containing TRIzol (1mL/mg; Life Sciences Technologies, Carlsbad, CA, USA), and RNA was extracted per manufacturers specifications. RNA concentration and purity (A260/280 ratios >1/8) were assessed using a Nanodrop 2000 (ThermoFisher Scientific) or BioTek Powerwave HT spectrophotometer (BioTek Instruments, Winooski, VT, USA). RNA (0.5-1 μg) was reverse-trancribed to cDNA using LunaScript RT SuperMix Kit (New England Biolabs, Ipswich, MA, USA). TaqMan gene expression assays with FAM-labeled probes were purchased from ThermoFisher Scientific (Carlsbad, CA, USA) to quantify expression of CYP1A1 (ThermoFisher Scientific; Hs01054797_g1), CYP1B1 (ThermoFisher Scientific; Hs00164383_m1), or AHRR (ThermoFisher Scientific; Hs01005075_m1). Real-time qPCR was performed on these targets using TaqMan Universal Master Mix (ThermoFisher Scientific) and a Quantstudio 3 Real-Time PCR system (Applied Biosystems, Waltham, MA, USA). Samples were run in triplicate and eukaryotic 18S (ThermoFisher Scientific; Hs03003631_g1) was used as an endogenous control. Relative gene expression was calculated using 2^-ΔΔCT^ from the respective control group.

### Statistics

The specific statistics used in each experiment are detailed in the legends for each figure. Generally, differences between groups were detected by Student’s t-test (2 groups/conditions), Analysis of Variance and a Sidak post-hoc test (3 or more groups/conditions), or Two-way Analysis of Variance and a Sidak post-hoc test (3 or more groups/conditions and two factors per group). P-values are stated in the legends for each figure, where P-values <0.05 were considered statistically significant. Values are presented as means ± Standard Deviation.

## Results

### Chronic Tobacco Smoke Exposure in Mice Adversely Affects Skeletal Muscle

To model an approximate smoking level of 1.5 packages of cigarettes per day over a 10-y period in humans, we exposed male mice to tobacco smoke for 60 min, 2x/d, 5d/week for 16 weeks (Fig. 1A). We observed lower body mass (Fig. 1B) and significant atrophy across a range of muscle phenotypes and function (Fig. 1C), accompanied by significant atrophy of individual muscle fibers (Fig. 1D). Muscle mounts to reveal neuromuscular junction morphology (Fig. 1E) in tibialis anterior (the fraction of abandoned endplates on this muscle has been previously published [17]) and diaphragm muscles revealed fragmentation of the acetylcholine receptor clusters and reduced motor axon diameter with chronic smoke exposure, and this was accompanied by a significant fraction of endplates lacking a detectable motoneuron (lack of synaptophysin; indicative of muscle fiber denervation) in both muscles (Fig. 1F). In addition, although chronic smoke exposure did not affect the protein abundance of the outer mitochondrial membrane protein voltage dependent anion channel, oxidative capacity was impaired as evident from a reduced maximal state III respiratory capacity in soleus and diaphragm muscles (Fig. 1G), suggesting the presence of an intrinsic mitochondrial oxidative defect in muscle with chronic smoke exposure. Next, we performed unbiased transcriptomics analysis (Fig. 2A-B) of skeletal muscle from control and smoke-exposed mice. Differential gene expression and network analysis found three major gene hubs showing robust reprogramming of the neuromuscular junction, myofibril/sarcomere, and the mitochondrion (Fig. 2C-E),

**Figure 1.**
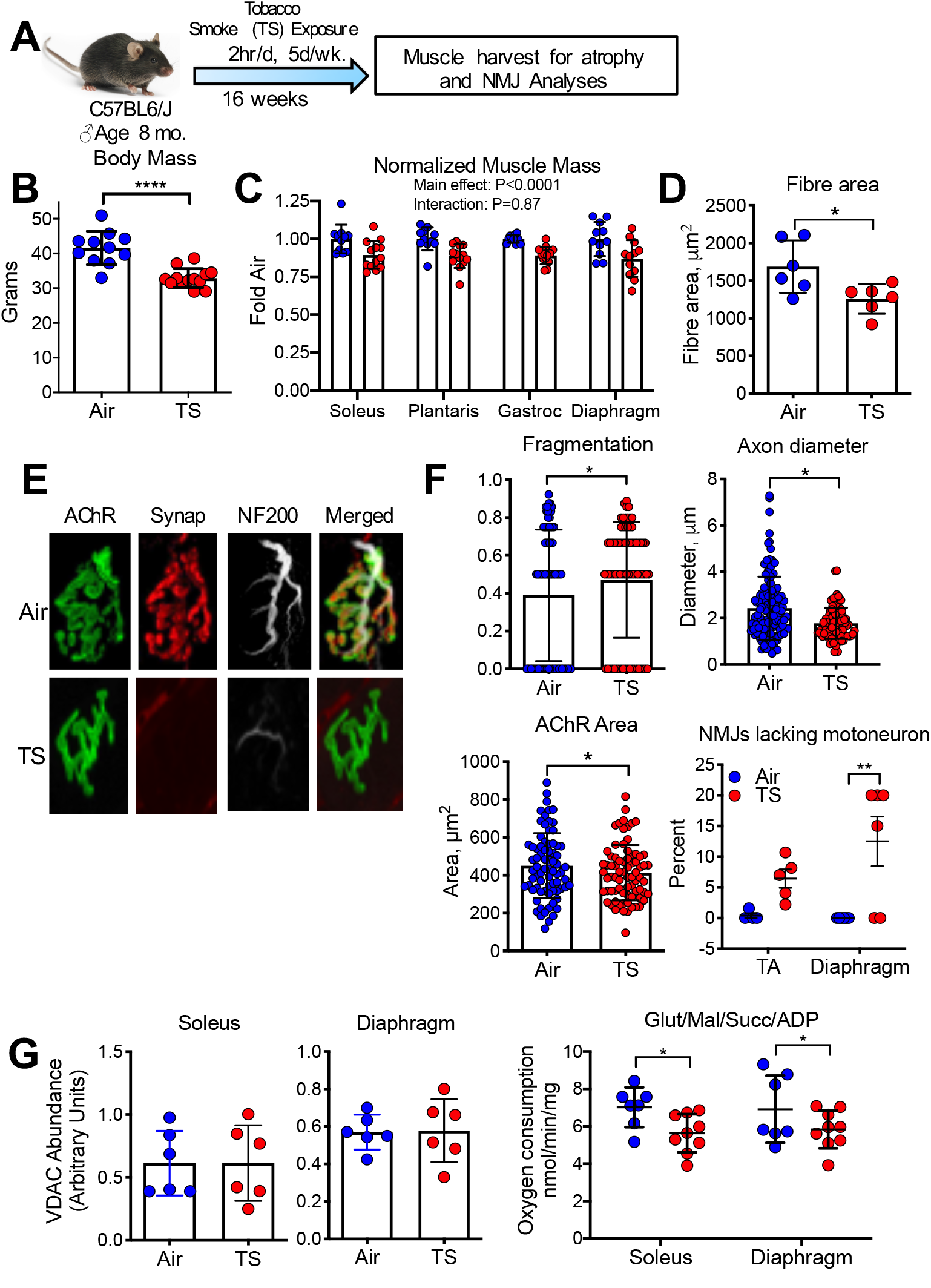
Male mice were exposed to mainstream TS for 60 min twice per d, 5 d per week, for 16 wk (A), and this caused a significant reduction in both body mass (B) (n = 11-13/group) and muscle mass across various muscles (C) (n = 11-13/group) and mean myofiber size in the soleus muscle (D) (n = 6/group). Neuromuscular junctions were labeled in TA muscle (AChRs, synaptophysin for motoneuron terminals) and Dia muscle (AChRs, synaptophysin, and NF200 for motor axons) and imaged by confocal microscopy (E). TS exposure caused AChR fragmentation (n = 75 NMJs from 6 mice/group), a reduction in axon diameter (n = 86-115 NMJs from 6 mice/group) and AChR area (n = 75 NMJs from 6 mice/group), and a significant accumulation of endplates that lacked detectable motoneuron terminals (lack of synaptophysin opposing AChRs) (n = 6 animals/group) (F). Although the mitochondrial outer membrane protein VDAC was no different between TS and Air exposed mice (n = 6/group), maximal ADP stimulated respiration with both complex I and II substrates was significantly reduced in soleus and diaphragm muscles from TS-exposed mice (G) (n = 7-9/group). Error bars = standard deviation. *P<0.05, **P<0.01, ****P<0.0001 using either two-tailed students *t*-test or ANOVA.

**Figure 2.**
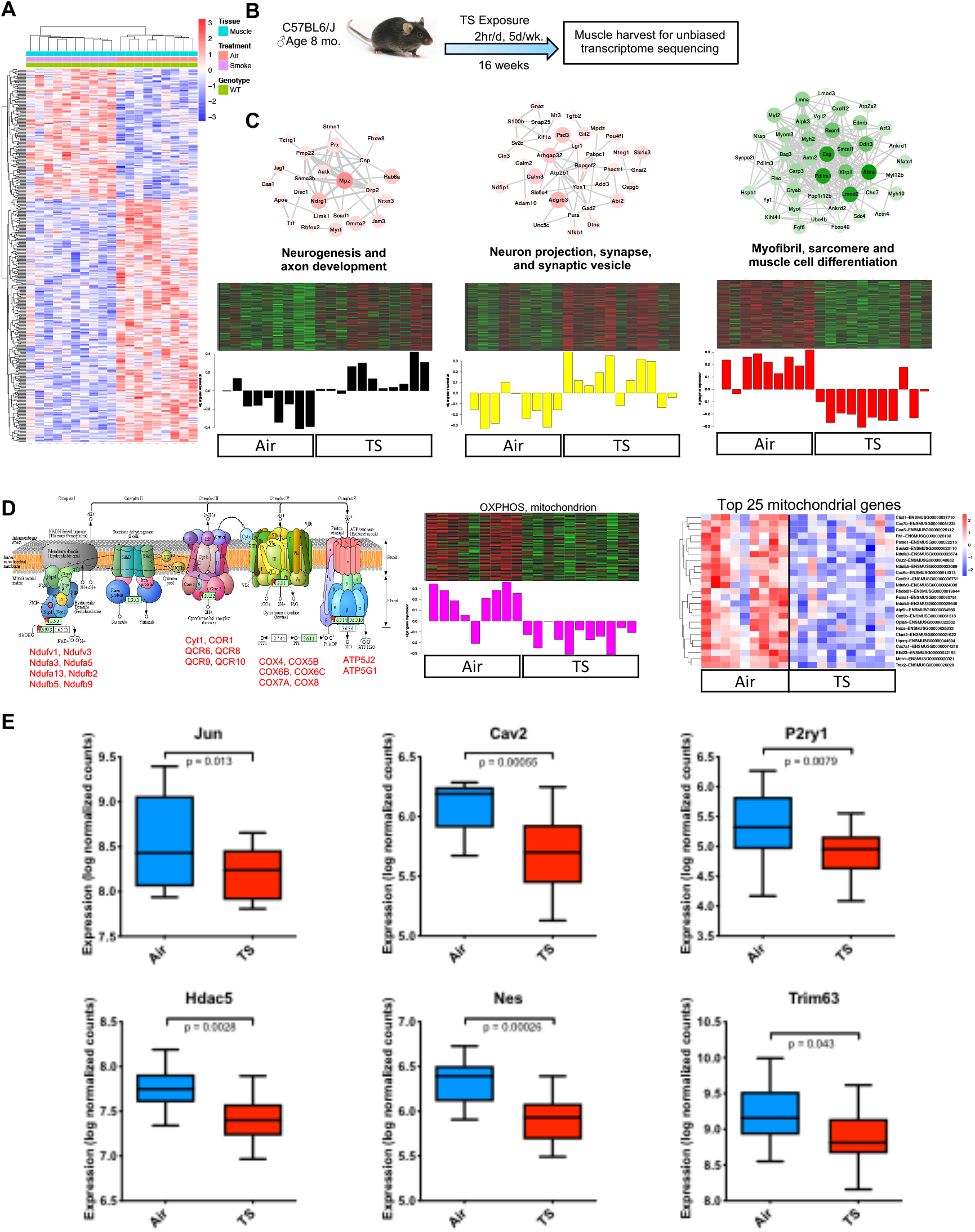
(A) Heatmap of differential gene expression analysis of RNAseq data from the plantaris muscle of mice exposed to (B) normal air or tobacco smoke for 16 weeks. The heatmap shows hierarchical clustering of 434 differentially expressed genes between air (n = 10) and TS-treated (n = 11) mice. (C) Network analysis of muscle gene expression profile changes identified significant ‘nodes’ corresponding to features related to the neuromuscular junction and myofibril. Node size (fold change) and color (p-value) (red = upregulation and green = downregulation) denotes extent of differential expression. Heatmaps depicting the expression of genes (rows) across samples (columns) for each node (red corresponds to gene upregulation and green to downregulation). (D) Additional significant gene nodes included oxidative phosphorylation and the mitochondrion with relevant heatmaps for all 140 genes in this node as well as the top 25 differentially expressed. (E) Boxplot representation of the expression levels of the genes associated with neuromuscular junction in air and smoke treated muscle samples. Non-parametric Wilcoxon test was used to compare differences between the datasets.

### Response of Skeletal Muscle AHR Signaling to Smoke Exposure

Various constituents of tobacco smoke have been shown to activate the AHR using a mouse hepatoma cell line [19]. To establish whether skeletal muscle responds to smoke exposure by up-regulating established AHR-regulated transcripts, we first determined the impact of 5 x 60 min acute smoke exposures performed over a 2.5-day period in mice on AHR signaling in skeletal muscle. This analysis revealed a trend (P=0.07) to an increase in the down-stream AHR effector Cyp1A1 with smoke exposure (Fig. 3A). Next, we examined the impact of smoking in human patients with two different diseases associated with chronic smoking and muscle atrophy: COPD [23] and critical limb ischemia (a severe form of peripheral artery disease) [22]. Consistent with observations in smoke-exposed mice, COPD patients that were current smokers had ~25-fold increased Cyp1A1 expression compared to COPD patients who were former smokers and compared to non-smoking control subjects (Fig. 3B). Similarly, muscle from active smoking critical limb ischemia patients had significant increases in the AHR-regulated transcripts Cyp1A1 and Cyp1B1, compared to non-smokers (Fig. 3C).

**Figure 3.**
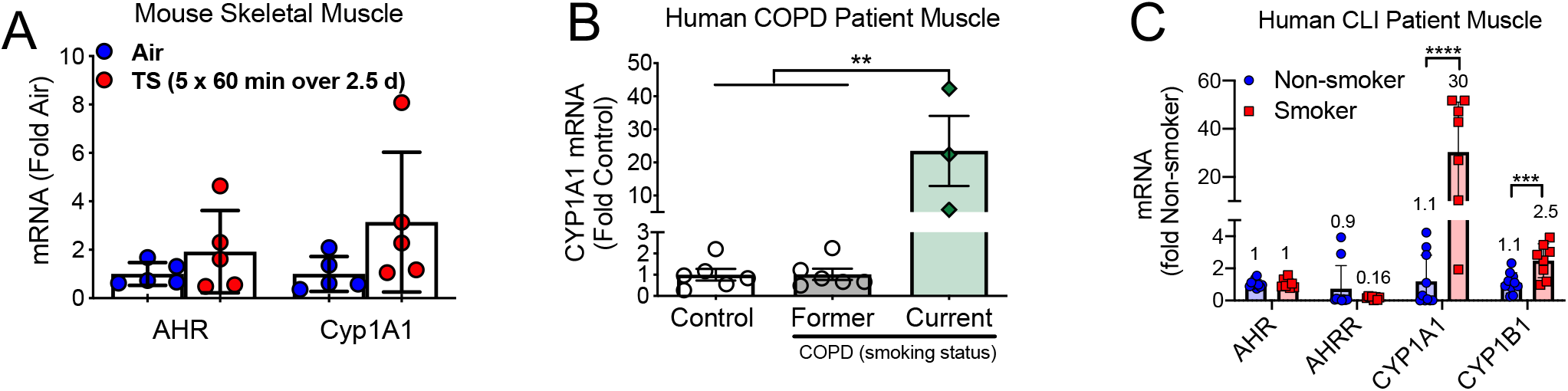
Smoking increases AHR signaling in mouse and human skeletal muscle. (A) Skeletal muscle from mice exposed to tobacco smoke (TS) exhibited elevated AHR and CYP1A1 mRNA expression (n = 5/group). (B) CYP1A1 mRNA expression in muscle biopsy specimens from human COPD patients and age-matched controls (n = 3-6/group). (C) AHR signaling is elevated in critical limb ischemia (CLI) patient muscle specimens obtained from smokers (n = 8-10/group). Statistical analysis performed by unpaired Student’s *t*-test or one-way ANOVA with Sidak post-hoc testing when necessary. **P<0.01, ***P<0.001, ****P<0.0001. Error bars = standard deviation.

### Tobacco Smoke Condensate Causes Atrophy and Impairs Mitochondrial Function

Using a C2C12 myotube culture system, we showed that treatment of mature myotubes with 0.02% tobacco smoke condensate did not cause appreciable cell death (Fig. 4A) but dramatically upregulated Cyp1A1 expression (confirming AHR activation) (Fig. 4B), and induced myotube atrophy (Fig. 4C). Smoke condensate also increased mitochondrial ROS emission (Fig. 4D) and reduced maximal oxidative capacity (Fig. 4E) in myotubes. We then showed that genetically antagonizing AHR signaling using a short hairpin RNA targeting the AHR (Fig. 4F) reduced Cyp1A1 expression in response to smoke condensate (Fig. 4G). Importantly, shAHR prevented the muscle atrophy induced by smoke condensate (Fig. 4H). A similar attenuation of smoke-induced Cyp1A1 expression (Fig. 4I) and myotube atrophy (Fig. 4J) was obtained when treating with the chemical AHR antagonists resveratrol [10] or CH223191 [18].

**Figure 4.**
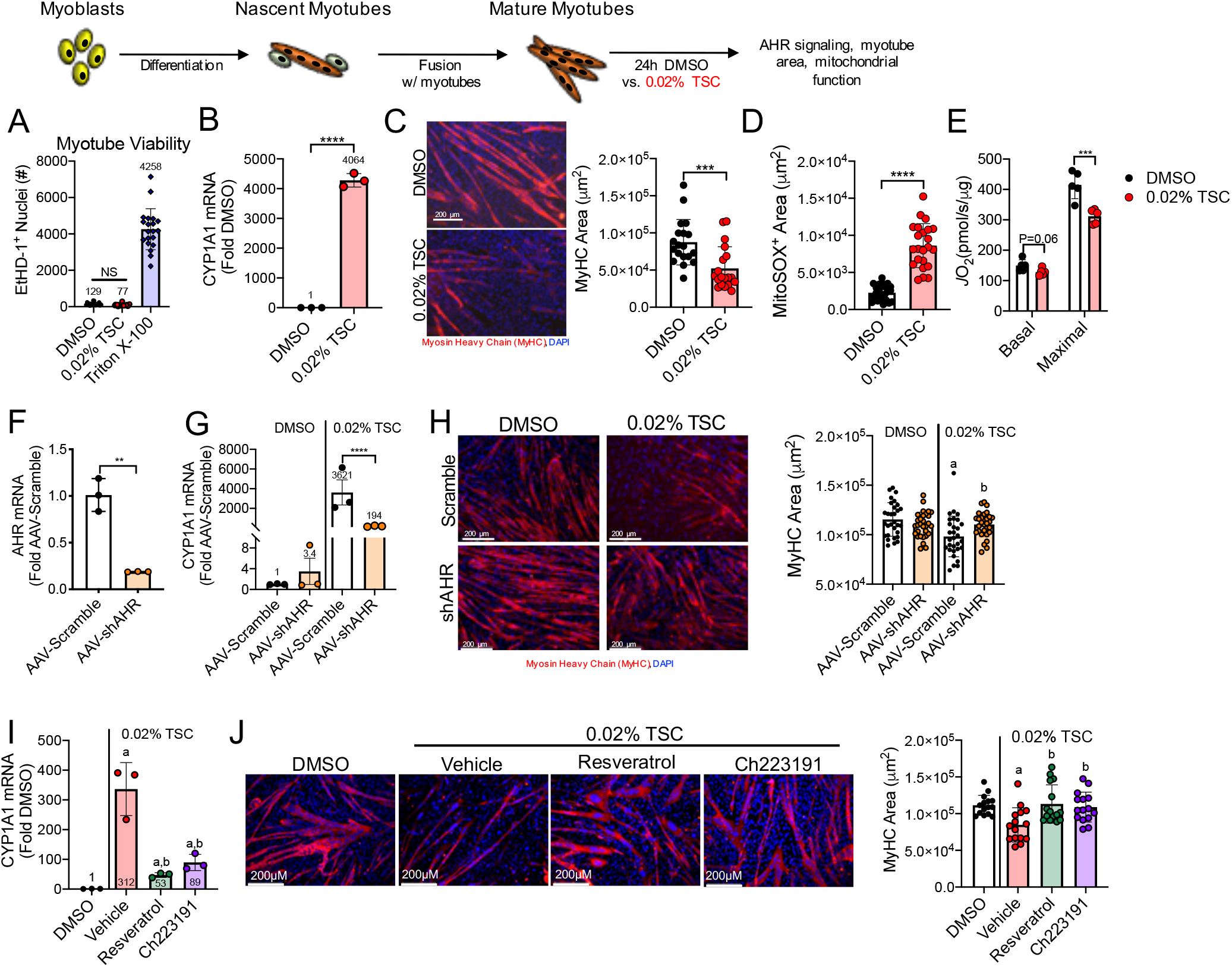
Tobacco smoke condensate causes myotube atrophy that is rescued by AHR antagonism. Treatment of C2C12 myotubes with tobacco smoke condensate (TSC) did not alter myotube viability (A) (n = 20/group), but resulted in increased CYP1A1 mRNA expression (B) (n = 3/group) and caused myotube atrophy (C) (n = 19/group), elevated mitochondrial ROS production (D) (n = 23/group), and impaired mitochondrial respiration (E) (n = 5/group). Expression of a short hairpin targeting the AHR (shAHR) decreased AHR mRNA expression (F) (n = 3/group) and attenuated CYP1A1 expression with TSC (G) (n = 3/group), and rescued myotube atrophy (H) (n = 30/group). Pharmacologic antagonism of the AHR with resveratrol (25μM) or CH223191 (1μM) decreased CYP1A1 mRNA (I) (n = 3/group) and rescued myotube atrophy (J) (n = 15/group). Statistical analysis performed by unpaired Student’s *t*-test or one-way ANOVA with Sidak post-hoc testing when necessary. **P<0.01, ***P<0.001, ****P<0.0001, ^a^P<0.05 vs. DMSO (within group), ^b^P<0.05 vs. between-group control (vehicle or AAV-Scramble). Error bars = standard deviation. NS = not significant.

### Chronic AHR Activity Without Smoke Exposure Causes Adverse Muscle Impact

To establish the impact of chronic AHR activity independent of tobacco smoke we constructed a mutant of the AHR that lacks the ligand binding domain and demonstrates constitutive activity (CAAHR; Fig. 5A), as done previously by another group [6]. Treatment of myotubes with AAV-CAAHR increased AHR expression 40-fold and increased Cyp1A1 10-fold relative to AAV-GFP treatment (Fig. 5B). Similar to observations with smoke condensate (Fig. 4), AAV-CAAHR treatment caused myotube atrophy (Fig. 5C), increased mitochondrial ROS, and impaired oxidative capacity (Fig. 5D).

**Figure 5.**
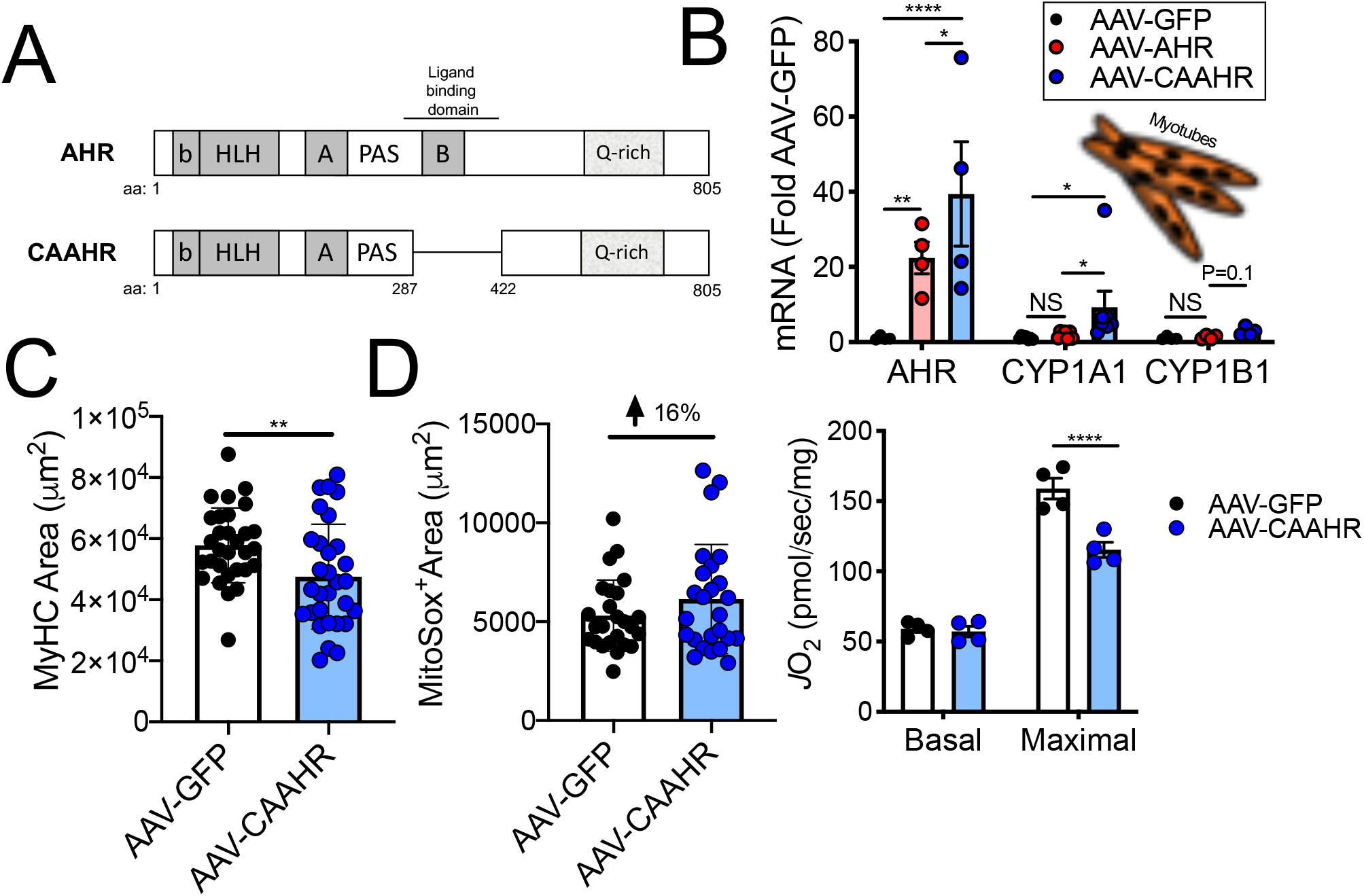
Viral-mediated expression of a constitutively active AHR causes myotube atrophy and mitochondrial impairments. (A) Simplified map showing the development of a constitutively active AHR (CAAHR) through deletion of the ligand binding domain. (B) AHR related mRNA signaling in myotubes (n = 4/group). (C) Quantification of myotube area (n = 30/group), and (D) mitochondrial ROS production (n = 25/group) and respiratory capacity with expression of AAV-GFP (control) or AAV-CAAHR (n = 4/group). Statistical analysis performed by unpaired Student’s *t*-test or one-way ANOVA with Sidak post-hoc testing when necessary. *P<0.05, **P<0.01, ****P<0.0001. Error bars = standard deviation. NS = not significant.

To establish the impact of chronic AHR activity independent of smoke exposure *in vivo*, we injected AAV containing the CAAHR mutant under the control of a cytomegalovirus promotor into one tibialis anterior muscle of C57BL6/J mice, and the contralateral tibialis anterior with AAV-Green Fluorescent Protein (Fig. 6A). AAV-CAAHR injection caused a >50-fold increase in AHR expression and a ~7-fold increase of Cyp1A1 (Fig. 6B). Similar to findings in AAV-CAAHR treated myotubes (Fig. 5C), and in mice chronically exposed to smoke (Fig. 1), muscle mass was reduced in the AAV-CAAHR injected limb in 3 out of 4 treated animals 12 weeks following AAV injection (Fig. 6C). In addition, AAV-CAAHR injection caused neuromuscular junction degeneration (Fig. 6D) that was characterized by acetylcholine receptor cluster fragmentation, reduced motor axon diameter, and a significant accumulation of endplates that lacked the motoneuron (Fig. 6E).

**Figure 6.**
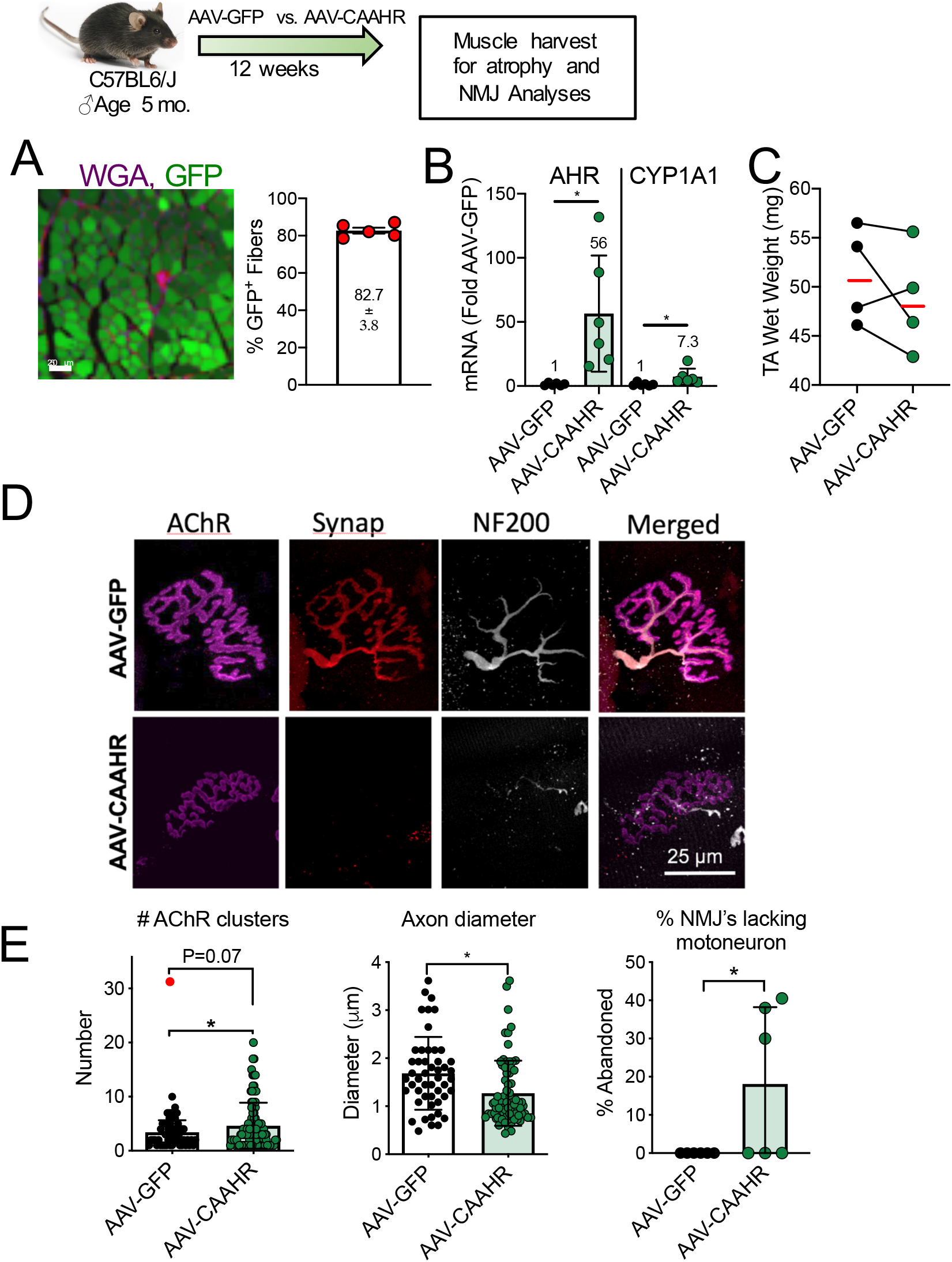
Viral-mediated expression of a constitutively active AHR in C57BL6/J mice causes muscle atrophy and NMJ degeneration. (A) Representative image and quantification of GFP+ myofibers of mouse TA muscle with AAV delivery (n = 5). (B) mRNA levels of AHR and CYP1A1 confirm constitutive AHR signaling in mouse muscles that received AAV-CAAHR (n = 5-6/group). (C) Wet weight of TA muscles from mice indicate smaller muscle size in AAV-CAAHR treated muscle (n = 4/group). (D) Representative images of NMJ morphology. (E) Quantitative analysis of NMJ morphology including acetylcholine receptor clusters (n = 60-100 NMJs from 5 mice/group), motor neuron axon diameters (n = 49-71 NMJs from 5 mice/group), and the percentage of NMJ’s lacking a motor neuron (indicative of denervation) (n = 6/group). Statistical analysis performed by unpaired Student’s *t*-test. *P<0.05. Error bars = standard deviation.

## Discussion

Long-term smoking is the primary cause of COPD and a significant fraction of patients develop muscle atrophy that predisposes them to worse clinical outcomes, including a greater risk of death [24]. Similarly, long-term smokers without disease also exhibit atrophy and reduced markers of mitochondrial oxidative capacity [21, 32], and chronic smoking in mouse models causes: muscle atrophy [9], reduced muscle oxidative capacity [11], and degeneration of the neuromuscular junction [16]. Despite this, the mechanisms by which smoking causes adverse muscle impact are poorly understood. In this respect, the AHR is a ligand-activated transcription factor responsive to a wide variety of environmental contaminants [27], including tobacco smoke [19], and chronic AHR activation can be toxic [28]. On this basis, we tested the hypothesis that chronic AHR activation, with or without tobacco smoke exposure, induces skeletal muscle atrophy, mitochondrial dysfunction and neuromuscular junction degeneration.

### Impact of Chronic TS Exposure on Skeletal Muscle

Based upon species-specific lifespan [3, 45], we modeled approximately a decade of 1-1.5 packs per day smoking behavior in humans by exposing mice to mainstream tobacco smoke for 2 h per d, 5 d per wk, for 16 wk. Smoke exposure in mice caused: muscle atrophy across a range of muscle phenotypes, muscle fiber atrophy, and reduced oxidative capacity. Furthermore, chronic smoke exposure induced pre- (reduced motor axon diameter, motoneuron terminal loss) and post-synaptic (acetylcholine receptor cluster fragmentation) neuromuscular junction alterations in both limb muscle and breathing muscle. Unbiased transcriptomics analysis supports these observations by showing that gene networks regulating mitochondria and the neuromuscular junction were amongst the most severely impacted with 16 wk smoke exposure. Furthermore, tobacco smoke condensate treatment of C2C12 myotubes induced a robust atrophy, elevated mitochondrial reactive oxygen species (higher mitoSox positive area) and impaired oxidative capacity. These observations are consistent with other studies suggesting that chronic smoking is a precipitating factor in the erosion of oxidative capacity [35, 36], muscle atrophy [46], and accumulation of denervated muscle fibers in muscle of COPD patients [17].

### Skeletal Muscle AHR Signaling with Smoke Exposure

Whilst adverse effects of smoking on skeletal muscle are relatively well-known and are attributed to a wide variety of chemicals within smoke [11], the mechanisms by which these chemicals adversely affect muscle are not well understood. The AHR is a ligand-activated transcription factor that regulates cytochrome P450 enzymes such as Cyp1A1 and Cyp1B1, as well as antioxidant pathways that include NAD[P]H quinone dehydrogenase and sulfiredoxin [27, 38]. Notably, the AHR has a wide variety of endogenous and exogenous ligands, including constituents of tobacco smoke [19]. Under normal (non-activated) conditions, most of the AHR pool is located in the cytoplasm and is bound with several chaperone proteins that prevent its nuclear translocation. However, upon ligand binding the chaperone proteins are released and permit the AHR to enter the nucleus where it binds to the AHR Nuclear Translocase. The AHR/AHR Nuclear Translocase dimer then binds to the so-called dioxin response element of target gene promotors to activate their transcription [27]. Whereas the AHR plays an important role in normal development [30], chronic AHR activation can have pathological consequences due to mitochondrial-mediated oxidative stress [1] for various organ systems [28], including the brain and nervous system, reproductive organs, heart, liver, and immune system [6]. To date, no studies have considered AHR-dependent toxicity in skeletal muscle, despite muscle atrophy being a unifying characteristic of conditions associated with chronic exposure to AHR ligands, including dioxin poisoning [25], chronic kidney disease [39], exposure to the herbicide Agent Orange [48], and long-term smoking [24].

To our knowledge, only three prior studies have mentioned the AHR in the context of skeletal muscle. One study showed that reduced AHR network gene expression was amongst the changes seen in a microarray of human muscle following resistance exercise training [34]. A second study detailed differential impact of exposure to the AHR agonist dioxin in skeletal muscle precursor cells depending upon their expression of the transcription factor Pax3 [12]. A third study was not focused on muscle *per se*, but used C2C12 muscle cells as a model to gain insight to the role of the AHR in cancer [5]. This last study showed that AHR activation in C2C12 muscle cells caused elevated mitochondrial reactive oxygen species and mitochondrial stress signaling in muscle cells [5], which is similar to other cell types where AHR activation increases mitochondrial reactive oxygen species generation and suppresses oxidative capacity [8, 40, 51]. Notably, our study is the first to address the impact of chronic tobacco smoke exposure on AHR signaling and its downstream consequences in skeletal muscle. In this respect, brief smoke exposure (5 repeated 60 min exposures over a 2.5 d period) in mice caused increased Cyp1A1 expression in limb muscle (P=0.07). Furthermore, our data from patients with diseases for which smoking is a key risk factor show a robust elevation of AHR signaling in skeletal muscle exclusively in current smokers. Thus, our data establish that the AHR pathway in skeletal muscle is responsive to tobacco smoke.

### Dependence of Smoke-induced Muscle Atrophy on AHR Signaling

Similar to *in vivo* observations in mouse and human skeletal muscle, treatment with tobacco smoke condensate in C2C12 myotubes *in vitro* robustly increased AHR signaling. Furthermore, smoke condensate exposure caused myotube atrophy without inducing cell death. Smoke condensate also increased mitochondrial reactive oxygen species emission and reduced oxidative capacity in myotubes. Not only did genetic antagonism using shRNA against the AHR attenuate Cyp1A1 expression with smoke condensate, it also prevented the smoke-induced atrophy. We further showed that the chemical AHR antagonists resveratrol [10] and CH223191 [18] also attenuated smoke-induced Cyp1A1 expression and myotube atrophy. These are the first data to establish that tobacco smoke exposure induces muscle atrophy in myotube culture and that this depends upon AHR signaling. As such, our results implicate an important role for smoke-induced AHR activation in the adverse muscle effects of chronic smoking.

### The Impact of Chronic AHR Activity in Skeletal Muscle Independent of Smoke Exposure

Previous studies have highlighted the toxicity that occurs with chronic AHR activation. This was first established in the context of dioxin poisoning where knockout of the AHR prevented much of the adverse responses to dioxin [13]. This was further advanced in studies showing that chronic AHR activity alone, induced by engineering mice to express a constitutively active AHR mutant, phenocopies many aspects of dioxin poisoning [6, 7]. However, these previous studies had not considered skeletal muscle. Thus, to establish the impact of chronic AHR activity in skeletal muscle, independent of tobacco smoke exposure, we constructed a mutant of the AHR that lacks the ligand binding domain and demonstrates constitutive activity, as has been done previously [6]. We then inserted this mutant into skeletal muscle using AAV in *in vitro* and *in vivo* models. In cultured C2C12 myotubes CAAHR transduction increased expression of both the AHR and its downstream target Cyp1A1, and, similar to tobacco smoke condensate, caused myotube atrophy and impaired mitochondrial function. Similar to these *in vitro* effects, 12 weeks following CAAHR transduction in tibialis anterior muscle of mice there was an increased expression of both the AHR and Cyp1A1 relative to the contralateral Green Fluorescent Protein transduced limb. Furthermore, CAAHR transduction reduced tibialis anterior muscle mass in 3 out of 4 animals relative to the contralateral limb, and induced remarkably similar alterations in pre-synaptic and post-synaptic structures of the neuromuscular junction to those seen with chronic smoke exposure. Thus, in multiple respects, our studies show that chronic AHR activity alone yields muscle changes that are very similar to the impact of chronic smoke exposure in mice.

### Potential Role of Chronic AHR Activity in Skeletal Muscle Impairment in COPD

The etiology of skeletal muscle impairment in COPD is multifactorial owing to the complexities of the disease pathophysiology (inflammation, oxidative stress, hypoxemia, hypercapnia, etc.) [24], and is also likely to include factors that precede disease onset. In this latter respect, long-term smoking is the most important proximate cause of COPD, and adverse effects of smoke exposure on muscle independent of disease are well-established [11, 21, 46]. Previous studies have identified a vast array of signaling pathways within skeletal muscle that respond to tobacco smoke exposure [33, 37, 43], but the AHR has not been considered previously. Indeed, although established AHR agonists including dioxins and polycyclic aromatic hydrocarbons are present in tobacco smoke [49], there may be thousands of chemicals at varying concentrations in tobacco smoke that are capable of activating the AHR [19]. On this basis, understanding the impact of chronic AHR activation in skeletal muscle is likely important to understanding the mechanisms involved in smoking-induced muscle impairment. In this respect, we show that smoke exposure robustly activates the AHR pathway in skeletal muscle, that this causes AHR-dependent atrophy, and that AHR activity in the absence of smoke exposure induces similar alterations in muscle as chronic smoke exposure. The significance of these observations is that they provide the first indication that chronic AHR activity induced by smoking is part of the complex etiology behind skeletal muscle impairment in COPD, and establish a rationale for preclinical therapeutic approaches targeting the AHR pathway.

## Conclusions

Muscle atrophy predisposes patients with TS-related diseases, such as COPD, to poor health outcomes that include greater mortality. To help us understand the mechanisms by which chronic smoking contributes to muscle impairment in COPD patients, the objective of our study was to test the hypothesis that chronic smoking-mediated activation of the AHR induces adverse muscle affect. Consistent with our hypothesis, we showed that tobacco smoke condensate caused myotube atrophy in an AHR-dependent manner. Similarly, knock-in of a constitutively active AHR mutant caused myotube atrophy and mitochondrial dysfunction. Finally, knock-in of a constitutively active AHR mutant in mouse muscle caused atrophy and produced similar changes in neuromuscular junction morphology as 16 wk of smoke exposure in a smoking mouse model. On the basis of our results, we suggest that chronic smoke-induced activation of the AHR plays an important role in the adverse muscle alterations seen in COPD patients that predispose them to exacerbated health outcomes.

